# Prenatal androgen treatment does not alter the firing activity of hypothalamic arcuate kisspeptin neurons in female mice

**DOI:** 10.1101/2021.07.19.452845

**Authors:** Amanda G. Gibson, Jennifer Jaime, Laura L. Burger, Suzanne M. Moenter

**Affiliations:** Departments of Molecular & Integrative Physiology, University of Michigan, Ann Arbor, MI, 48109, USA; Departments of Neuroscience Graduate Program, University of Michigan, Ann Arbor, MI, 48109, USA; Departments of Internal Medicine, University of Michigan, Ann Arbor, MI, 48109, USA; Departments of Obstetrics & Gynecology, University of Michigan, Ann Arbor, MI, 48109, USA; Departments of Reproductive Sciences Program, University of Michigan, Ann Arbor, MI, 48109, USA

## Abstract

Neuroendocrine control of reproduction is disrupted in many individuals with polycystic ovary syndrome, who present with increased luteinizing hormone (LH), and presumably gonadotropin-releasing hormone (GnRH), release frequency, and high androgen levels. Prenatal androgenization (PNA) recapitulates these phenotypes in primates and rodents. Female offspring of mice injected with dihydrotestosterone (DHT) on gestational D16-18 exhibit disrupted estrous cyclicity, increased LH and testosterone, and increased GnRH neuron firing rate as adults. PNA also alters the developmental trajectory of GnRH neuron firing rates, markedly blunting the prepubertal peak in firing that occurs in 3wk-old controls. GnRH neurons do not express detectable androgen receptors and are thus probably not the direct target of DHT. Rather, PNA likely alters GnRH neuronal activity by modulating upstream neurons, such as hypothalamic arcuate neurons co-expressing kisspeptin, neurokinin B (gene Tac2), and dynorphin, aka KNDy neurons. We hypothesized PNA treatment changes firing rates of KNDy neurons in a similar age-dependent manner as GnRH neurons. We conducted targeted extracellular recordings (0.5-2h) of Tac2-identified KNDy neurons from control and PNA mice at 3wks of age and in adulthood. About half of neurons were quiescent (<0.005Hz). Long-term firing rates of active cells varied, suggestive of episodic activity, but were not different among groups. Short-term burst firing was also similar. We thus reject the hypothesis that PNA alters the firing rate of KNDy neurons. This does not preclude altered neurosecretory output of KNDy neurons, involvement of other neuronal populations, or *in-vivo* networks as critical drivers of altered GnRH firing rates in PNA mice.

**Significance statement:** Prenatal androgenization (PNA) recapitulates key aspects of the common reproductive disorder polycystic ovary syndrome. It is postulated that disruptions in the episodic pattern of gonadotropin-releasing hormone (GnRH) secretion in part underly this disorder, yet GnRH neurons do not express androgen receptor to respond directly to elevated androgens. A population of kisspeptin, neurokinin B, and dynorphin-expressing (KNDy) neurons in the hypothalamic arcuate nucleus are thought to regulate pulsatile GnRH release and some express androgen receptor. We did not find evidence, however, that PNA altered spontaneous activity of KNDy neurons before puberty at 3wks of age or in adulthood. This suggests that PNA likely acts through other components of the broader hypothalamic network to change the patterns of GnRH release.

## Introduction

Gonadotropin-releasing hormone (GnRH) regulates the secretion of the gonadotropins luteinizing hormone (LH) and follicle-stimulating hormone (FSH) from the anterior pituitary. GnRH is released in an episodic manner that varies in frequency through the female reproductive cycle (Levine and Ramirez, 1982; Moenter et al., 1991). Lower frequency favors synthesis and release of FSH over LH and is important for recruiting ovarian follicles and their subsequent maturation, whereas higher pulse frequency in the mid-late follicular phase favors LH, which drives androgen synthesis (Haisenleder et al., 1991; Wildt et al., 1981). Failure to vary the frequency of GnRH release is thought to be a key neuroendocrine phenotype of the reproductive disorder polycystic ovary syndrome (PCOS).

PCOS is a complex spectrum of reproductive and metabolic phenotypes with postulated genetic and environmental causes. Patients with PCOS often exhibit a persistently high LH (and presumably GnRH) pulse frequency, leading to disrupted follicle maturation and ovulation. Increased LH stimulation also drives hyperandrogenism (Dumesic et al., 2015; McCartney et al., 2002). To investigate the etiology of the disorder, animal models are needed for experimental manipulations and measurements that cannot be conducted in humans. Elevated prenatal androgen exposure (PNA) recapitulates PCOS-like reproductive phenotypes in many species including non-human primates (Abbott et al., 2008, 2017, 2019; Dumesic et al., 1997), sheep (Birch et al., 2003; Veiga-Lopez et al., 2008), and rodents (Foecking et al., 2005; Sullivan and Moenter, 2004). In adult PNA mice, LH pulse frequency (Moore et al., 2015) and GnRH-neuron action potential firing frequency (Roland and Moenter, 2011) are both increased. PNA in mice also alters the developmental trajectory of GnRH-neuron firing frequency, which is interesting as aspects of PCOS may emerge around the pubertal transition (McCartney et al., 2002; Rosenfield, 2007). Specifically, in control mice, the firing frequency peaks at 3wks of age before decreasing to adult levels (Dulka and Moenter, 2017). In contrast, the firing frequency in PNA female mice did not vary with age, and it was lower than control mice at 3wks, distinct from the increase observed in PNA adults (Dulka and Moenter, 2017; Sullivan and Moenter, 2004)

The mechanisms by which PNA alters the activity of GnRH neurons are not completely understood. These neurons do not express detectable levels of androgen receptor (Herbison et al., 1996), thus it is likely that upstream neuronal populations are involved in regulating their firing patterns. One such population is in the hypothalamic arcuate nucleus, specifically neurons that co-express kisspeptin, neurokinin B, and dynorphin (KNDy neurons). KNDy neurons are posited to be involved in the control of pulsatile GnRH, and subsequent LH, secretion (Clarkson et al., 2017; S. Y. Han et al., 2015; McQuillan et al., 2019). KNDy neurons express receptors for gonadal steroids, including androgen receptor (Smith et al., 2005), and could serve as the site of steroidal feedback that alters GnRH neuron activity and/or a site of action for PNA exposure (Caldwell et al., 2017; Oakley et al., 2009; Vanacker et al., 2017; Walters et al., 2018).

We hypothesized that PNA treatment would alter the firing frequency of KNDy neurons in an age-dependent manner similar to that of GnRH neurons. We tested this by assessing the effect of PNA on the spontaneous firing frequency of KNDy neurons in prepubertal 3wk-old and adult female mice through long-term extracellular recordings. Specifically, we postulated that PNA treatment would increase KNDy-neuron activity relative to controls in adults but reduce activity relative to controls in 3-wk old mice. We also predicted that 3wk-old control mice would exhibit increased KNDy-neuron firing frequency relative to control adults and 3wk-old PNA mice.

## Materials and Methods

### Animals

Mice expressing enhanced green fluorescent protein (GFP) under the control of Tac2 promoter (Tac2-GFP, BAC transgenic mice (015495-UCD/STOCK Tg [Tac2-EGFP]381Gsat, Mouse Mutant Regional Resource Center (http://www.mmrrc.org/) were used to identify KNDy neurons for recording. In mice, *Tac2* encodes neurokinin B, which is co-expressed with kisspeptin and dynorphin in KNDy neurons. Tac2-GFP-identified cells in brain slices used for recording also express kisspeptin and/or dynorphin at high percentages, supporting their identity as KNDy neurons (Ruka et al., 2013). Mice were maintained in a 14h:10h dark photoperiod (lights on at 0300 Eastern Standard Time) and had *ad libitum* access to water and either Harlan 2919 chow during pregnancy/lactation or 2916 chow for maintenance. All animal procedures were performed in accordance with the University of Michigan Institutional Animal Care and Use Committee’s regulations.

To generate experimental mice, a Tac2-GFP female and CD1 female mice were bred with a C57B/6 male and monitored daily for a copulatory plug (day 1 of pregnancy). The CD1 dam assists in providing maternal care and nutrition. On days 16-18 of pregnancy, dams were injected subcutaneously with 225μg/day of dihydrotestosterone (DHT) or sesame oil as vehicle. Control offspring from dams for whom timing of pregnancy could not be clearly established were also included in studies without injections; firing rate from these mice did not differ from vehicle-treated mice (treatment: *F*_(1, 22)_ = 2.114, *p* = 0.160; interaction of age and treatment: F_(1, 22)_ = 0.237, p = 0.631). Experiments were conducted on female offspring prior to weaning at 3 weeks

of age (PND 18-21) or in adulthood (PND 66-152; median 133). PNA status was confirmed by anogenital distance and estrous cyclicity in adults. Anogenital distance was measured with digital calipers on 2-3 successive days and averaged for each mouse. Estrous cyclicity was assessed via vaginal cytology and studies on adult females were done on diestrus. Cycle stage was confirmed with uterine mass; one DHT-treated mouse was excluded due to a uterine mass of 136.2mg, suggestive of incorrect cycle identification based on vaginal cytology.

### Brain Slice Preparation

All solutions were bubbled with 95% O_2_/5% CO_2_ for at least 15min prior to tissue exposure and throughout the procedures. The brain was rapidly removed and cooled for 60s in ice-cold sucrose saline solution containing (in mM): 250 sucrose, 3.5 KCL, 26 NaHCO_3_, 10 D-glucose, 1.25 Na_2_HPO_4_, 1.2 MgSO4, and 3.8 MgCl_2_. Coronal slices (300 µm) through the hypothalamic region, including the ARC, were cut with a Leica VT1200S (Leica Biosystems, Buffalo Grove, IL). Slices were incubated for 30min at room temperature in 50% sucrose saline and 50% artificial cerebral spinal fluid (ACSF) containing (in mM): 135 NaCl, 3.5 KCl, 26 NaHCO_3_, 10 D-glucose, 1.25 Na_2_HPO_4_, 1.2 MgSO_4_, and 2.5 CaCl_2_ (pH 7.4). The slices were then held in 100% ACSF at room temperature for between 0.5 to 5.5h before recording. No differences in results were attributable to duration post brain slice preparation.

### Electrophysiological recording

To evaluate the long-term firing patterns of KNDy neurons with minimal disruption of the cell’s intrinsic properties, targeted single-unit extracellular recordings were conducted (Alcami et al., 2012; Nunemaker et al., 2003). Individual slices were transferred to a recording chamber mounted on the stage of an Olympus BX51WI upright fluorescent microscope. A constant perfusion of ACSF at a rate of 3mL/min was established with a MINIPULS 3 peristaltic pump (Gilson, Middleton, WI). The chamber was maintained at a temperature of 29-32°C with an inline heating system (Warner Instrument Corporation, Hamden, CT). ACSF was replaced every hour.

Recording electrodes (resistance 2-4MΩ) were pulled from borosilicate glass (Schott no. 8250; World Precision Instruments, Sarasota, Fl) using a Sutter P-97 puller (Sutter Instrument, Novato, CA). The pipettes were filled with a HEPES-buffered pipette solution containing (in mM): 150 NaCl, 10 HEPES, 10 D-glucose, 2.5 CaCl_2_, 1.3 MgCl_2_, and 3.5 KCl, pH7.4. At the surface of the brain slice, a small amount of negative pressure was applied to bring the pipette in contact with tissue, facilitating the later formation of a low-resistance seal (<100MΩ) between the pipette and neuron (Alcami et al., 2012). Recordings were made with one channel of an EPC10 dual patch-clamp amplifier using PatchMaster software (HEKA Elektronic, Lambrecht,Germany). Cells were held in voltage-clamp with a 0mV pipette holding potential. Seal resistance was checked every 10-15min by measuring response to a 5mV hyperpolarizing step between series. Data were acquired at 10kHz and filtered at 5kHz.

Recording duration ranged from 0.5 to 2.6h (mean±SEM 71.7±2.9min; median 60min). If a cell was not firing at the conclusion of a recording session, either high-potassium ACSF (20 mM K^+^) or the neurokinin 3 receptor agonist senktide (100nM; Phoenix Pharmaceuticals, Burlingame, CA) was bath-applied. If a cell failed to respond to these stimuli with increased firing, recording integrity could not be verified and data analysis was truncated to the last action current.

Response to senktide was quantified by comparing the spontaneous firing frequency for the 5min prior to addition of senktide to the ACSF to the firing frequency for 5min, beginning 2min after senktide reached the bath to allow time for the drug to equilibrate in the chamber and penetrate the slice. This 2min delay was chosen based on the onset of and peak senktide response across cells.

### Analysis

Event detection was completed using IgorPro8 (WaveMetrics, Inc., Lake Oswego, OR) utilizing custom routines, and all events were manually confirmed. The average spontaneous firing rate was calculated for each cell as total events/recording duration. The short-term patterns of neuronal activity were also assessed with custom IgorPro8 routines. Repetitive, grouped firing events are referred to as “bursts” for analysis. To be considered part of a burst, a firing event must occur within a defined “burst window” after the previous event. The burst window for analysis of these KNDy neurons was identified by varying the burst window from 0.01s to 1s in 10ms intervals and selecting the burst window that captures the maximal burst frequency for the control cells; this was 230ms as in prior reports (Vanacker et al., 2017). At the selected burst window, the software characterizes each event as belonging to a burst or as a single spike, then calculates the following parameters: burst frequency, burst duration, intraburst interval, spikes per burst, single spike frequency, and interevent interval. Burst duration and spikes per burst are the averages for all bursts from a given cell. Intraburst interval is the average of intervals between spikes in a burst, whereas interevent interval is the average of intervals greater than the burst window and can occur between bursts, between single spikes or between single spikes and bursts. Short breaks in the recording (typically <2s) occur at 10-15 min intervals to monitor the seal resistance. Intervals that crossed these gaps were not included when calculating cells’ averages. Spikes that occurred within 230ms (i.e., the burst window) of these gaps, or the start or end of the recording, were characterized according to the available information; this could lead to an underestimate of the burst frequency, burst duration, and/or spikes per burst.

### PCR to assess arcuate gene expression

Hypothalamic tissue punches were collected to assess the effect of PNA on gene expression. Separate cohorts of mice from those used for recordings were utilized to collect tissue micro-punches from the ARC. A coronal slice was obtained with an adult mouse brain matrix (1mm, Zivic Instruments, Pittsburgh, PA); an initial cut was made just caudal to the optic chiasm, followed by a cut just rostral to the brain stem (2-3 mm thick) for the ARC. Tissue punches were made with a 1.2mm Palkovits punch. Tissue was immediately homogenized in RLT buffer (Qiagen, Valencia, California) containing 2-mercaptoethanol (1%v/v, Sigma), snap frozen, and stored at -80°C. RNA from was extracted with the RNeasy Micro Kit with on column DNasing (Qiagen). 240ng RNA per sample was reverse transcribed with Superscript IV VILO Master Mix (Fisher/Invitrogen). A standard curve of hypothalamic RNA (600, 120, 24, 4.8 and 0ng/20µl) was also reverse transcribed (Ruka et al., 2013). The transcripts for: *Kiss1, Kiss1r, Pdyn, Oprk1, Tac2, Tacr3, Ar, Esr1 and Pgr* were assayed via Taqman quantitative PCR in duplicate with10ng cDNA. Data were analyzed by the ΔΔCT method (Bustin, 2002), normalized to *Actb* and *Syn1*and reported relative to 3wk-CON. Primers and Taqman probes were purchased from Integrated DNA Technologies (Coralville, IA) and are reported in Table 1.

**Table 1.**
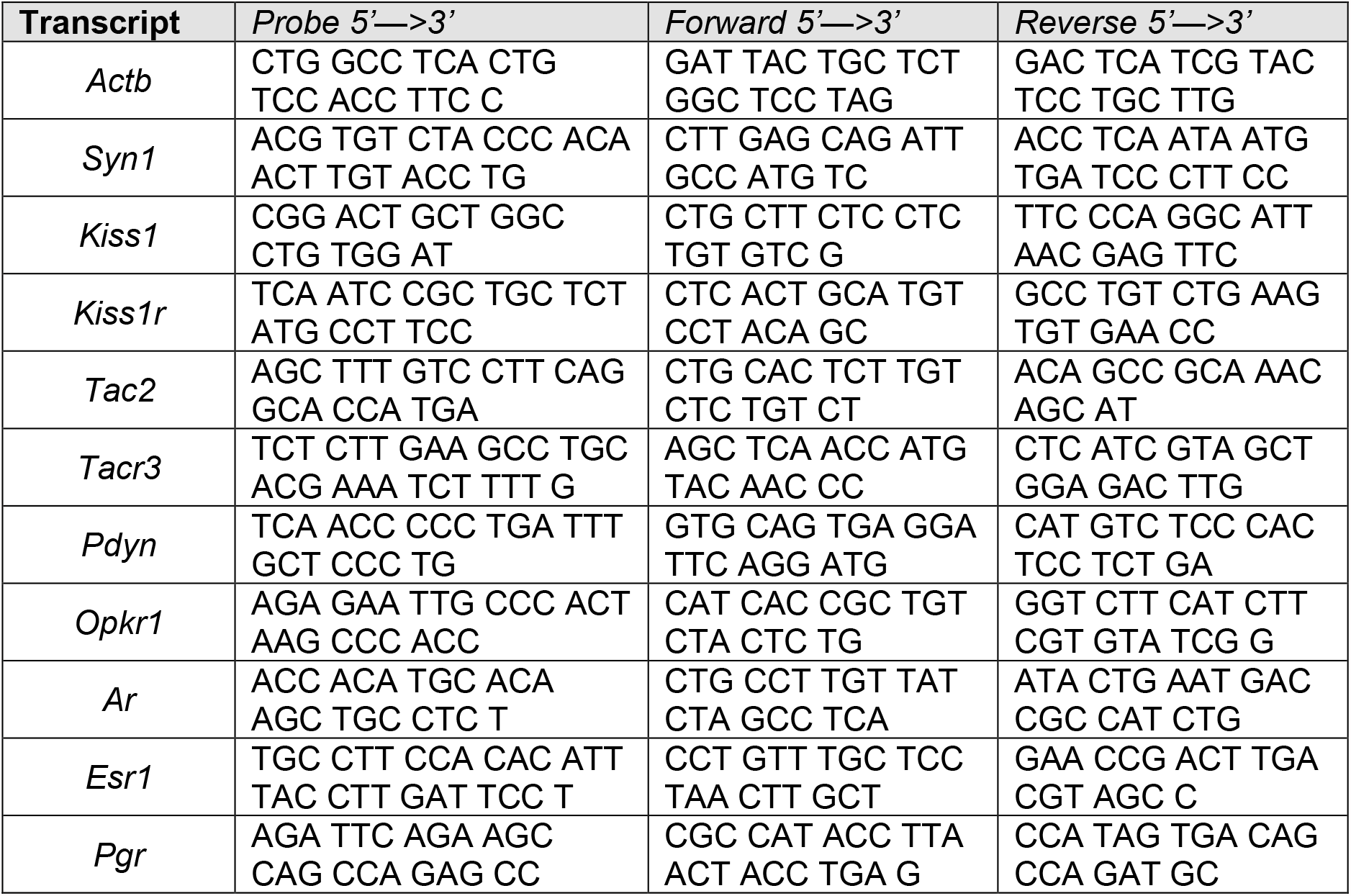
Probes and primers sequences for PCR experiments (Figure 5).

### Statistics

Data visualization and analyses were conducted with R (R Core Team, 2019) and RStudio (RStudio Team, 2019) using a combination of open-sourced packages (Chang, 2014; Chang et al., 2020; Fox and Weisberg, 2019; Gohel, 2020b, 2020a; Henry and Wickham, 2020b, 2020a; Kassambara, 2020, 2021; Schauberger and Walker, 2020; Schloerke et al., 2020; Wickham et al., 2019, 2020; Wickham and Hester, 2020; Wickham and Seidel, 2020; Wilke, 2020; Xie, 2014, 2015, 2020; Xie et al., 2021; Zhu, 2020) and custom procedures. Additional statistical analyses were conducted with Prism 9 (GraphPad, La Jolla, CA). Data are reported as mean±SEM, with median illustrated where indicated. For recordings, n is number of cells; for PNA phenotype confirmation and mRNA quantification, n is number of mice. Normality of the data distribution was evaluated with Shapiro-Wilk. Two-way ANOVA (Type III) was conducted to evaluate the main effects and interactions of age and prenatal treatment.

Bonferroni correction for multiple comparisons was used as test is sufficiently robust for non-normally distributed data (Underwood, 1997). The level accepted as significant was set to *p*<0.05. Statistical tables for two-way ANOVAs report the differences in means and associated 95% CI defined for age (adult – 3wk), treatment (PNA – control), and interaction ([adult PNA – adult control] – [3wk PNA – 3wk control]).

### Software accessibility

The event detection and burst analysis code described in the paper is freely available online at https://gitlab.com/um-mip/coding-project. The R analysis code is freely available online at https://github.com/gibson-amandag/PNA_KNDy. The code is available as Extended Data 1. Analyses were conducted on a MacBook Pro, Early 2015 version, running macOS Catalina 10.15.7 and on a Mac Mini, 2018 version, running macOS Mojave 10.14.6

## Results

### PNA characterization

To verify the effects of PNA, anogenital distance, body mass, and estrous cycles were recorded from adult mice and the surviving female littermates of 3wk-old mice where possible. Adult PNA mice had a longer anogenital distance (Figure 1A, statistical parameters in Table 2, control n=17 mice from 10 litters, PNA 23 mice from 11 litters, *p*<0.0001) and larger body mass (Figure 1B, p=0.026) than control mice. PNA treatment also altered the distribution of days spent in each estrous cycle stage (Figure 1C, D, p<0.0001). PNA mice spent more days in diestrus than expected (standardized residual=7.08) and fewer days in proestrus than expected (standardized residual=-10.11). These results indicate that the PNA treatment was successful.

**Table 2.**
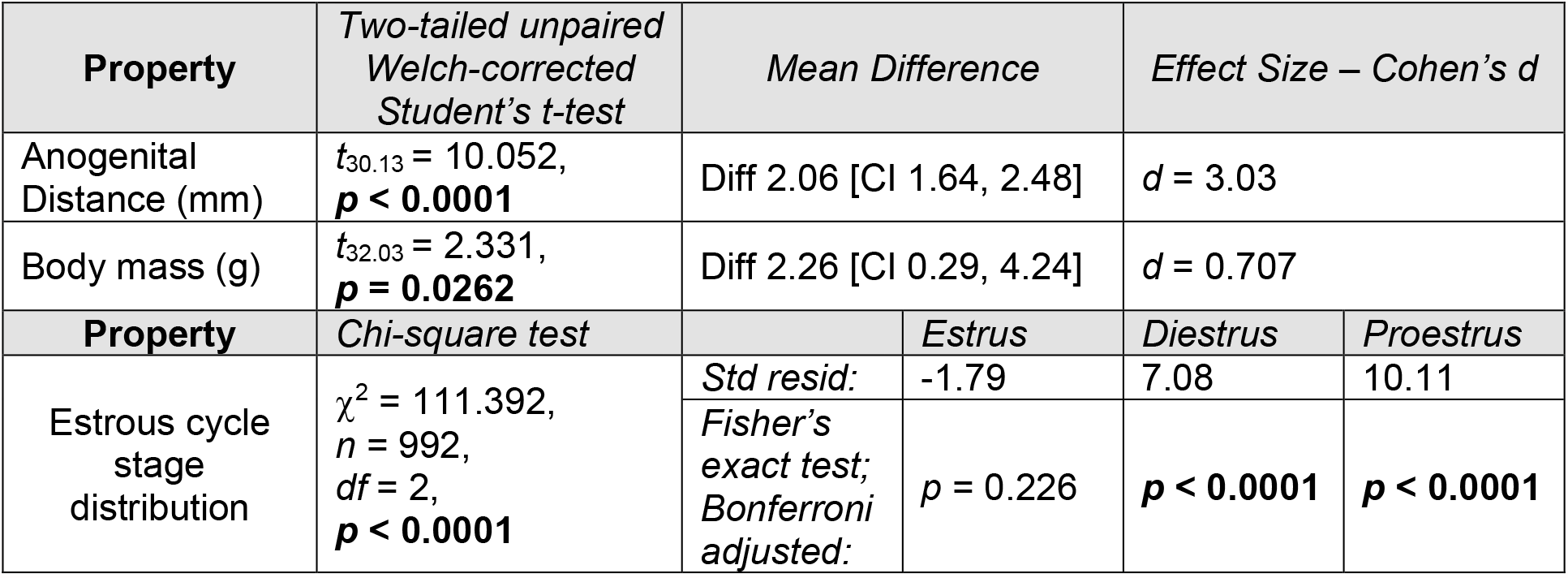
Statistical parameters characterizing the PNA phenotype (Figure 1). Bold indicates *p*<0.05. Std resid, standardized residuals of PNA group from Chi-square test

| Property | Two-tailed unpaired<br>Welch-corrected<br>Student's *t*-test | Mean Difference |  | Effect Size – Cohen's *d* |  |  |
| --- | --- | --- | --- | --- | --- | --- |
| Anogenital<br>Distance (mm) | $t_{30.13} = 10.052$,<br>**$p < 0.0001$** | Diff 2.06 [CI 1.64, 2.48] | | $d = 3.03$ | | |
| Body mass (g) | $t_{32.03} = 2.331$,<br>**$p = 0.0262$** | Diff 2.26 [CI 0.29, 4.24] | | $d = 0.707$ | | |
| Property | Chi-square test |  | Estrus | Diestrus | Proestrus |  |
| Estrous cycle<br>stage<br>distribution | $\chi^2 = 111.392$,<br>$n = 992$,<br>$df = 2$,<br>**$p < 0.0001$** | Std resid: | -1.79 | 7.08 | 10.11 | |
| | | Fisher's<br>exact test;<br>Bonferroni<br>adjusted: | $p = 0.226$ | **$p < 0.0001$** | **$p < 0.0001$** | |

**Figure 1.**
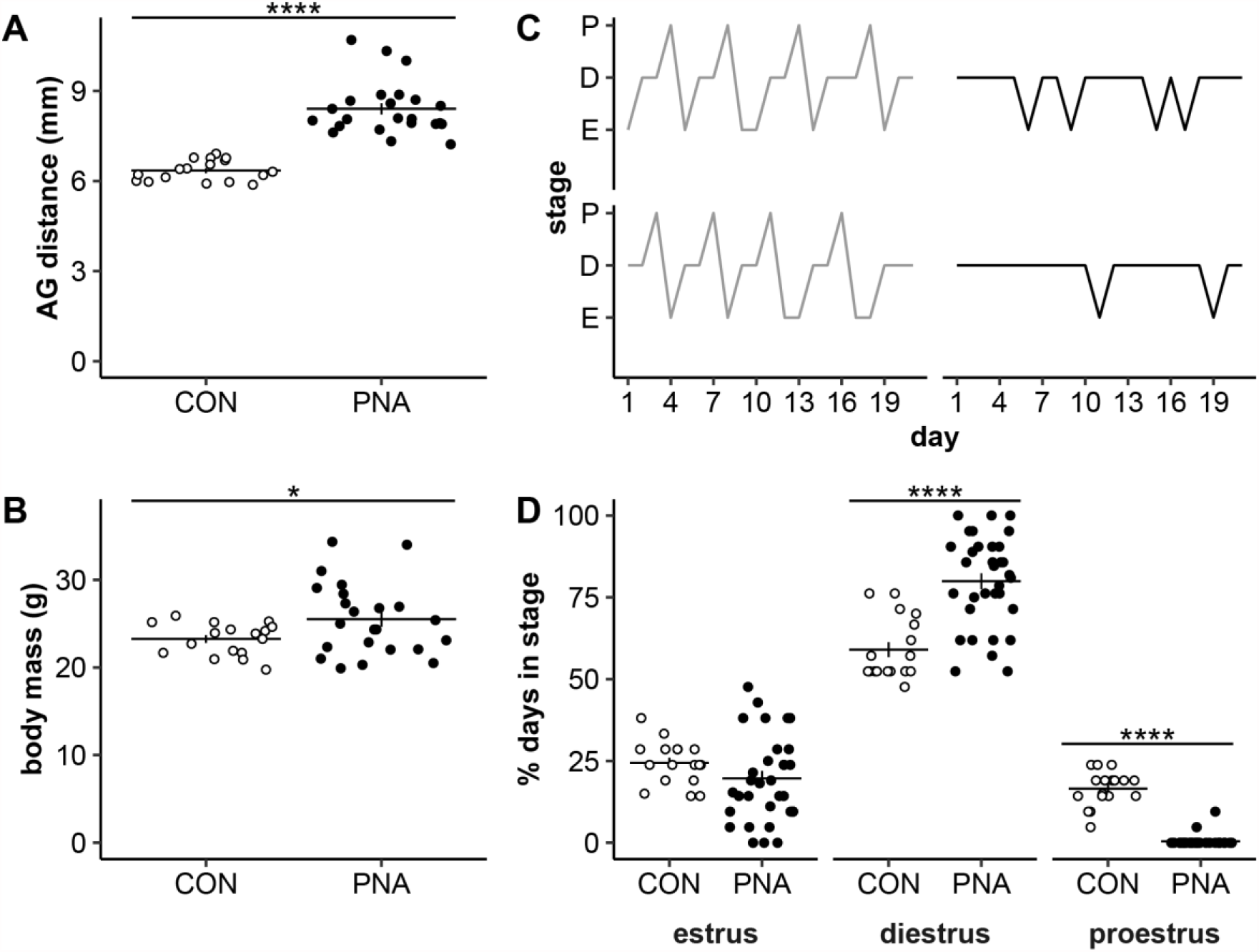
Confirmation of PNA phenotype in adults and surviving littermates of 3wk mice used for electrophysiology. A, B, D. Individual values (CON open symbols, PNA black symbols) and mean ± SEM anogenital (AG) distance (A), body mass (B) or percent days in each cycle stage (D). C. Representative estrous cycles over a 3-week period; P proestrus, D diestrus, E estrus. Statistical parameters reported in Table 2. *p<0.05, ****p≤0.0001

### Spontaneous firing rate

To determine how age and PNA treatment alter the firing activity of KNDy neurons, we conducted targeted, long-term extracellular recordings of Tac2-GFP-identified neurons in the arcuate nucleus of the hypothalamus. These neurons exhibited firing patterns consistent with episodic activity (representative traces in Figure 2A and 2B). About half of recorded Tac2-GFP neurons were quiescent (defined as <0.005Hz). The proportion of quiescent neurons did not vary with age or treatment (Figure 2C; statistical parameters reported in Table 3; 3wk-CON n=11 cells from 7 mice in 5 litters, 3wk-PNA n=22 cells from 12 mice in 6 litters, adult-CON n= 15 cells from 10 mice in 6 litters, adult-PNA n=22 cells from 13 mice in 8 litters). Neither age nor PNA treatment affected the mean firing frequency of Tac2-GFP neurons over the recording period (Figure 2D).

**Table 3.** Statistical parameters for firing activity (Figure 2). Independence of treatment and age with firing proportion was assessed with Breslow-Day test. This was followed by the Mantel-Haenszel Chi-squared test with continuity correction to determine the effect of treatment on firing proportion when controlling for age (Simonoff, 2003). A two-way ANOVA was conducted for firing frequency.

| ***Parameter*** | ***Breslow-Day test for independence*** |  | ***Mantel-Haenszel Chi-squared*** |
| --- | --- | --- | --- |
| Proportion firing | $\chi^2 = 1.285, df = 1, p = 0.257$ | | $\chi^2 = 0.069, df = 1, p\text{-value} = 0.7932$ |
| ***Parameter*** | ***age*** | ***PNA treatment*** | ***interaction*** |
| firing frequency | Diff -0.193<br>[CI -0.537, 0.151] | Diff -0.104<br>[CI -0.448, 0.240] | Diff 0.130<br>[CI -0.558, 0.819] |
| | $F(1, 66) = 1.251;$<br>$p = 0.267$ | $F(1, 66) = 0.362;$<br>$p = 0.549$ | $F(1, 66) = 0.143;$<br>$p = 0.707$ |

**Figure 2.**
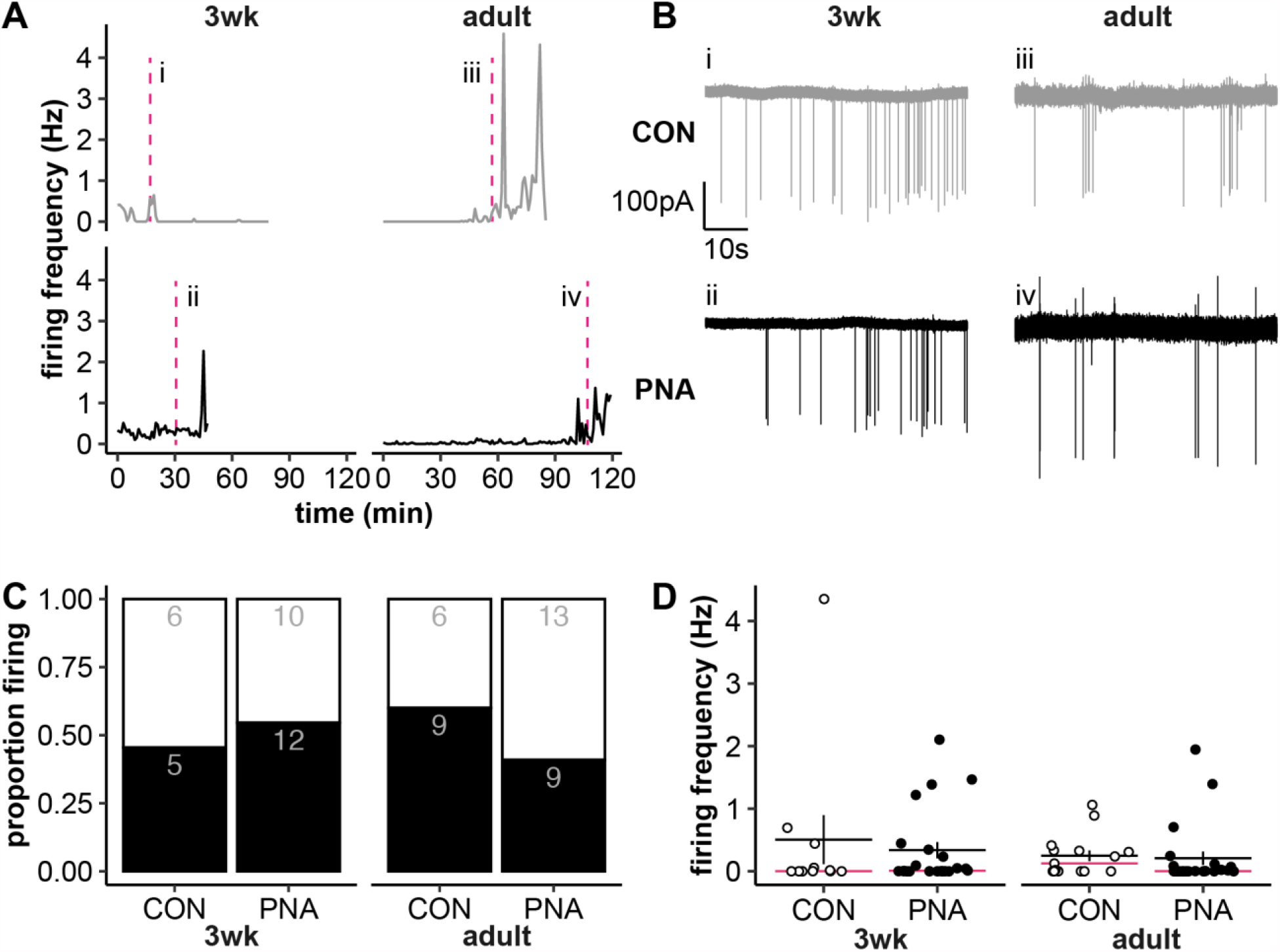
Effect of age and PNA treatment on KNDy neuron firing activity. A, Representative long-term firing patterns (1-min bins). CON are shown on the top in grey, PNA on the bottom in black for each age. The time of the traces shown in B is designated by the magenta dashed lines in panel A. B. Examples of raw firing data (60s) from the areas indicated in A, details as in A. The selected 60s bins are representative of the mean firing rate of each group. C. Proportion of cells with a firing frequency >0.005Hz (black bars) vs <0.005Hz (white bars); numbers are cell counts in each group. D. Individual values (CON open symbols, PNA black symbols) and mean ± SEM and median (magenta line) for firing frequency across the duration of long-term recordings. Statistical parameters reported in Table 3.

### Response to Senktide

To verify viability of quiescent cells, the neurokinin B receptor agonist senktide was added at the conclusion of a subset of spontaneous recordings (3wk-CON n=8 cells from 6 mice in 5 litters, 3wk-PNA n=12 cells from 8 mice in 4 litters, adult-CON n=8 cells from 6 mice in 3 litters, adult-PNA n=14 cells from 10 mice in 6 litters). While performed as a quality check, this test is also biologically relevant as senktide activates firing activity of Tac2-GFP neurons (Ruka et al., 2013), and it is possible that age and treatment alter this. Tac2-GFP neurons responded to senktide with an increase in firing frequency (Figure 3; statistical parameters in Table 4, main effect of time, p<0.0001). This increase was evident in adult-CON (*p*=0.007), adult-PNA (*p*=0.001), and 3wk-CON (*p*=0.003), yet there was not a significant increase for the 3wk-PNA (*p*=0.415). This suggests that PNA treatment may alter the development of the response to senktide in KNDy neurons.

**Table 4.** Statistical parameters from three-way mixed model ANOVA assessing the effect of age and PNA treatment on the KNDy cell response to senktide (Figure 3). The effect size, generalized *η*^*2*^, is reported for each effect. Bonferroni multiple comparisons for cells that differ by only one factor. Bold indicates *p* < 0.05.

| Effect | Statistic | generalized $\eta^2$ |
| --- | --- | --- |
| senktide | $F(1, 38) = 52.35, p < 0.0001$ | 0.395 |
| age | $F(1, 38) = 0.7515, p = 0.391$ | 0.010 |
| PNA treatment | $F(1, 38) = 2.235, p = 0.143$ | 0.030 |
| senktide x age | $F(1, 38) = 0.5459, p = 0.465$ | 0.007 |
| senktide x PNA treatment | $F(1, 38) = 2.051, p = 0.160$ | 0.025 |
| age x treatment | $F(1, 38) = 0.2633, p = 0.611$ | 0.004 |
| senktide x age x PNA treatment | $F(1, 38) = 1.179, p = 0.284$ | 0.015 |
| **Bonferroni's multiple comparisons test** | *Mean diff (Hz), 95% CI of diff* | *Statistic* |
| SK 3wk-CON – C 3wk-CON | Diff 2.36, [CI 0.59, 4.13] | $t_{38} = 4.055, \mathbf{p = 0.003}$ |
| SK 3wk-PNA – C 3wk-PNA | Diff 1.04, [CI -0.41, 2.49] | $t_{38} = 2.192, p = 0.415$ |
| SK adult-CON – C adult-CON | Diff 2.178, [CI 0.40, 3.95] | $t_{38} = 3.743, \mathbf{p = 0.007}$ |
| SK adult-PNA – C adult-PNA | Diff 2.00, [CI 0.66, 3.34] | $t_{38} = 4.540, \mathbf{p = 0.001}$ |
| C 3wk-PNA – C 3wk-CON | Diff 0.11, [CI -1.51, 1.72] | $t_{76} = 0.194, p > 0.999$ |
| C adult-PNA – C adult-CON | Diff -0.18, [CI -1.75, 1.39] | $t_{76} = 0.340, p > 0.999$ |
| SK 3wk-PNA – SK 3wk-CON | Diff -1.21, [CI -2.82, 0.40] | $t_{76} = 2.223, p = 0.350$ |
| SK adult-PNA – SK adult-CON | Diff -0.36, [CI -1.93, 1.20] | $t_{76} = 0.682, p > 0.999$ |
| C adult-CON – C 3wk-CON | Diff 0.19, [CI -1.58, 1.95] | $t_{76} = 0.315, p > 0.999$ |
| C adult-PNA – C 3wk-PNA | Diff -0.10, [CI -1.49, 1.29] | $t_{76} = 0.207, p > 0.999$ |
| SK adult-CON – SK 3wk-CON | Diff 0.01, [CI -1.76, 1.77] | $t_{76} = 0.011, p > 0.999$ |
| SK adult-PNA – SK 3wk-PNA | Diff 0.86, CI [-0.53, 2.25] | $t_{76} = 1.825, p = 0.863$ |

**Figure 3.**
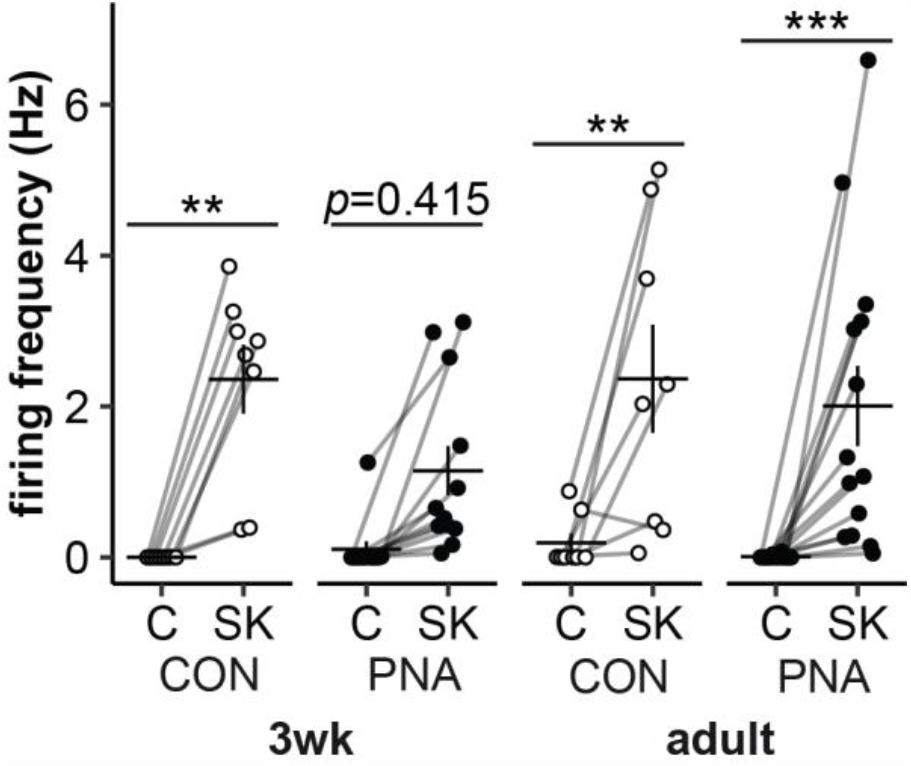
Senktide activates KNDy-neuron firing activity. Individual values and mean ± SEM of firing rate during 5min control (C) and senktide (SK) periods. CON mice are shown in open symbols, PNA mice in black symbols. ** p≤0.01, *** p≤0.001. Statistical parameters in Table 4.

### Short-term firing pattern

Examining the average firing frequency over the duration of the recording could obscure changes in the short-term organization of action potentials that may be more relevant for neurosecretion (Cazalis et al., 1985; Dutton and Dyball, 1979). We thus investigated the effect of age and PNA treatment on short-term firing patterns called bursts. (Figure 4; statistical parameters in Table 5). Because not all cells exhibit burst firing, the n for cells changes for parts B, C and D, and for part F as detailed in the legend. There were no differences due to age or treatment on any parameter other than burst duration. Burst duration was greater in cells from adults than those from 3wk-old mice (Figure 4B; *p*=0.031). An increase in burst duration could occur as a result of more spikes per burst, and/or a longer intraburst interval. Though it did not reach the level set for statistical significance, the increase in burst duration in adults appears to be driven primarily by increased spikes per burst (Figure 4C; *p*=0.096) rather than a change in the intraburst interval (Figure 4D; *p*=0.911).

**Table 5.** Two-way ANOVA statistical parameters for burst parameters. Bold indicates *p*<0.05.

| Property | age | PNA treatment | interaction |
| --- | --- | --- | --- |
| burst frequency | Diff -0.030<br>[CI -0.086, 0.025] | Diff -0.014<br>[CI -0.070, 0.041] | Diff 0.023<br>[CI -0.088, 0.134] |
| | $F(1, 66) = 1.184$;<br>$p = 0.281$ | $F(1, 66) = 0.267$;<br>$p = 0.608$ | $F(1, 66) = 0.167$;<br>$p = 0.684$ |
| burst duration | Diff 0.116<br>[CI 0.011, 0.221] | Diff 0.063<br>[CI -0.042, 0.168] | Diff 0.116<br>[CI -0.94, 0.326] |
| | $F(1, 35) = 5.068$;<br>**$p = 0.031$** | $F(1, 35) = 1.479$;<br>$p = 0.232$ | $F(1, 35) = 1.263$;<br>$p = 0.269$ |
| spikes per burst | Diff 0.954<br>[CI -0.178, 2.087] | Diff 0.560<br>[CI -0.572, 1.693] | Diff 0.955<br>[CI -1.310, 3.220] |
| | $F(1, 35) = 2.927$;<br>$p = 0.096$ | $F(1, 35) = 1.009$;<br>$p = 0.322$ | $F(1, 35) = 0.733$;<br>$p = 0.398$ |
| intraburst interval | Diff 0.002<br>[CI -0.039, 0.044] | Diff 0.021<br>[CI -0.021, 0.062] | Diff -0.003<br>[CI -0.085, 0.079] |
| | $F(1, 35) = 0.013$;<br>$p = 0.911$ | $F(1, 35) = 1.021$;<br>$p = 0.319$ | $F(1, 35) = 0.005$;<br>$p = 0.942$ |
| single spike frequency | Diff -0.073<br>[CI -0.187, 0.041] | Diff -0.015<br>[CI -0.129, 0.099] | Diff -0.122<br>[CI -0.350, 0.106] |
| | $F(1, 66) = 1.63$; $p = 0.206$ | $F(1, 66) = 0.072$;<br>$p = 0.790$ | $F(1, 66) = 1.147$;<br>$p = 0.288$ |
| interburst interval | Diff -14.87<br>[CI -84.09, 54.36] | Diff 13.72<br>[CI -55.51, 82.94] | Diff 87.73<br>[CI -50.71, 226.2] |
| | $F(1, 42) = 0.188$;<br>$p = 0.667$ | $F(1, 42) = 0.160$;<br>$p = 0.691$ | $F(1, 42) = 1.635$;<br>$p = 0.208$ |

**Figure 4.**
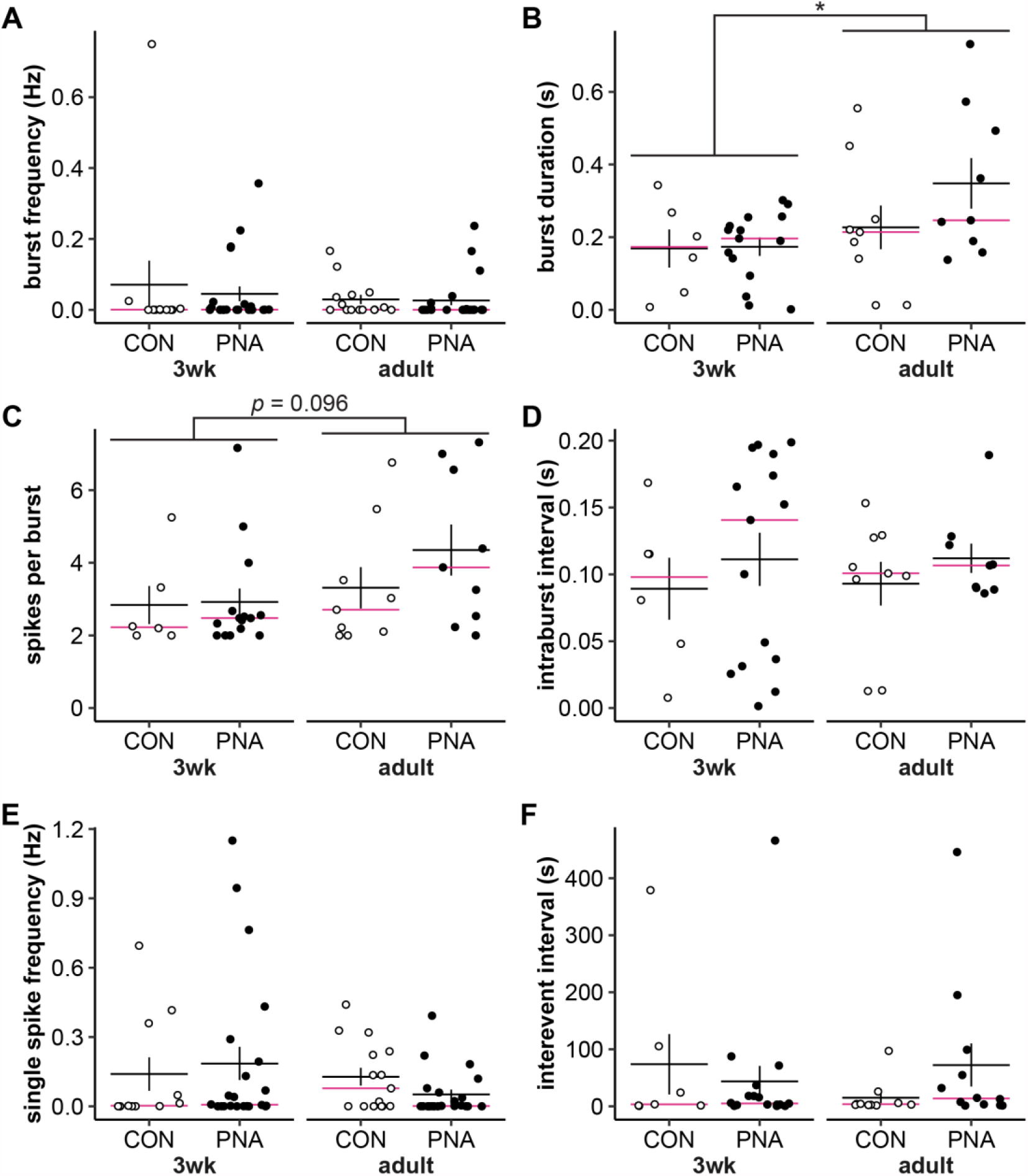
Effect of age and PNA treatment on burst parameters. A-F. Individual values (CON open symbols, PNA black symbols) and mean ± SEM and median (magenta lines) burst frequency (A), burst duration (B), spikes per burst (C), intraburst interval (D), single spike frequency (E), and interevent interval (F). B, C, D. Burst parameters are only calculated for cells with at least one burst; 3wk-CON n=6 cells from 5 mice in 4 litters, 3wk-PNA n=15 cells from 10 mice in 5 litters, adult-CON n=9 cells from 8 mice in 5 litters, adult-PNA n=9 cells from 8 mice in 5 litters. F. Calculating interevent interval also requires multiple events, 3wk-CON n=7 cells from 6 mice in 5 litters, 3wk-PNA n=17 cells from 10 mice in 5 litters, adult-CON n=10 cells from 10 mice in 6 litters, adult-PNA n=12 cells from 9 mice in 5 litters; *p<0.05. Statistical parameters in Table 5.

### Development but not PNA affects expression of key transcripts

To examine the effects of age and PNA on steroid receptors and KNDy neuron peptides and receptors in the arcuate nucleus, we quantified mRNA expression of androgen (*Ar*), estrogen (*Esr1*), and progesterone (*Pgr*) receptors and of kisspeptin (*Kiss1*), neurokinin B (*Tac2*), and dynorphin (*Pdyn*), and their corresponding receptors (*Kiss1r, Tacr3*, and *Oprk1*, respectively). *Tac2* (p=0.0001) and *Tacr3* (p<0.0001) expression were both increased in adults compared to 3wk-old mice (Figure 5, statistical parameters in Table 6, 3wk-CON n=9 mice, 3wk-PNA n=8 mice, adult-CON n=8 mice, adult-PNA n=7 mice). Similarly, *Ar* (p<0.0001) and *Pgr* (p<0.0001) were increased in the adult arcuate nucleus (Figure 5). PNA treatment did not alter expression of any transcripts, though there were weak trends for PNA to increase expression of *Kiss1r* (p=0.091) and *Pdyn* (p=0.071, Figure 5).

**Table 6.** Two-way ANOVA statistical parameters for arcuate mRNA expression. Bold indicates *p*<0.05.

| Transcript | age | PNA treatment | interaction |
| --- | --- | --- | --- |
| *Actb* | Diff -0.018<br>[CI -0.074, 0.038]<br>$F(1, 28) = 0.436$;<br>$p = 0.515$ | Diff -0.019<br>[CI -0.074, 0.037]<br>$F(1, 28) = 0.466$;<br>$p = 0.501$ | Diff -0.019<br>[CI -0.131, 0.092]<br>$F(1, 28) = 0.126$;<br>$p = 0.725$ |
| *Syn1* | Diff 0.018<br>[CI -0.038, 0.074]<br>$F(1, 28) = 0.436$;<br>$p = 0.515$ | Diff 0.019<br>[CI -0.037, 0.074]<br>$F(1, 28) = 0.466$;<br>$p = 0.501$ | Diff 0.019<br>[CI -0.092, 0.131]<br>$F(1, 28) = 0.126$;<br>$p = 0.725$ |
| *Kiss1* | Diff 0.068<br>[CI -0.358, 0.495]<br>$F(1, 28) = 0.108$;<br>$p = 0.744$ | Diff 0.019<br>[CI -0.407, 0.445]<br>$F(1, 28) = 0.009$;<br>$p = 0.926$ | Diff 0.028<br>[CI -0.824, 0.880]<br>$F(1, 28) = 0.004$;<br>$p = 0.947$ |
| *Kiss1r* | Diff -0.019<br>[CI -0.170, 0.132]<br>$F(1, 28) = 0.068$;<br>$p = 0.796$ | Diff 0.129<br>[CI -0.022, 0.280]<br>$F(1, 28) = 3.062$;<br>$p = 0.091$ | Diff 0.150<br>[CI -0.152, 0.453]<br>$F(1, 28) = 1.038$;<br>$p = 0.317$ |
| *Tac2* | Diff 1.075<br>[CI 0.584, 1.566]<br>$F(1, 28) = 20.113$;<br>**$p = 0.0001$** | Diff 0.009<br>[CI -0.482, 0.500]<br>$F(1, 28) = 0.001$;<br>$p = 0.97$ | Diff 0.264<br>[CI -0.718, 1.245]<br>$F(1, 28) = 0.302$;<br>$p = 0.587$ |
| *Tacr3* | Diff 1.064<br>[CI 0.707, 1.420]<br>$F(1, 28) = 37.321$;<br>**$p < 0.0001$** | Diff 0.108<br>[CI -0.249, 0.464]<br>$F(1, 28) = 0.381$;<br>$p = 0.542$ | Diff -0.476<br>[CI -1.189, 0.238]<br>$F(1, 28) = 1.865$;<br>$p = 0.183$ |
| *Pdyn* | Diff 0.138<br>[CI -0.224, 0.499]<br>$F(1, 28) = 0.608$;<br>$p = 0.442$ | Diff 0.331<br>[CI -0.030, 0.693]<br>$F(1, 28) = 3.520$;<br>$p = 0.071$ | Diff -0.153<br>[CI -0.876, 0.571]<br>$F(1, 28) = 0.187$;<br>$p = 0.669$ |
| *Opkr1* | Diff 0.127<br>[CI -0.044, 0.298]<br>$F(1, 28) = 2.313$;<br>$p = 0.139$ | Diff 0.095<br>[CI -0.076, 0.266]<br>$F(1, 28) = 1.294$;<br>$p = 0.265$ | Diff -0.170<br>[CI -0.512, 0.172]<br>$F(1, 28) = 1.033$;<br>$p = 0.318$ |
| *Ar* | Diff 0.673<br>[CI 0.493, 0.854]<br>$F(1, 28) = 58.323$;<br>**$p < 0.0001$** | Diff 0.041<br>[CI -0.140, 0.221]<br>$F(1, 28) = 0.215$;<br>$p = 0.647$ | Diff 0.150<br>[CI -0.212, 0.511]<br>$F(1, 28) = 0.720$;<br>$p = 0.403$ |
| *Esr1* | Diff 0.091<br>[CI -0.195, 0.377]<br>$F(1, 28) = 0.428$;<br>$p = 0.519$ | Diff 0.002<br>[CI -0.284, 0.288]<br>$F(1, 28) = 0.0002$;<br>$p = 0.989$ | Diff -0.105<br>[CI -0.677, 0.467]<br>$F(1, 28) = 0.141$;<br>$p = 0.71$ |
| *Pgr* | Diff 0.955<br>[CI 0.585, 1.325]<br>$F(1, 28) = 27.92$;<br>**$p < 0.0001$** | Diff -0.114<br>[CI -0.484, 0.257]<br>$F(1, 28) = 0.395$;<br>$p = 0.535$ | Diff 0.076<br>[CI -0.664, 0.817]<br>$F(1, 28) = 0.045$;<br>$p = 0.834$ |

**Figure 5.**
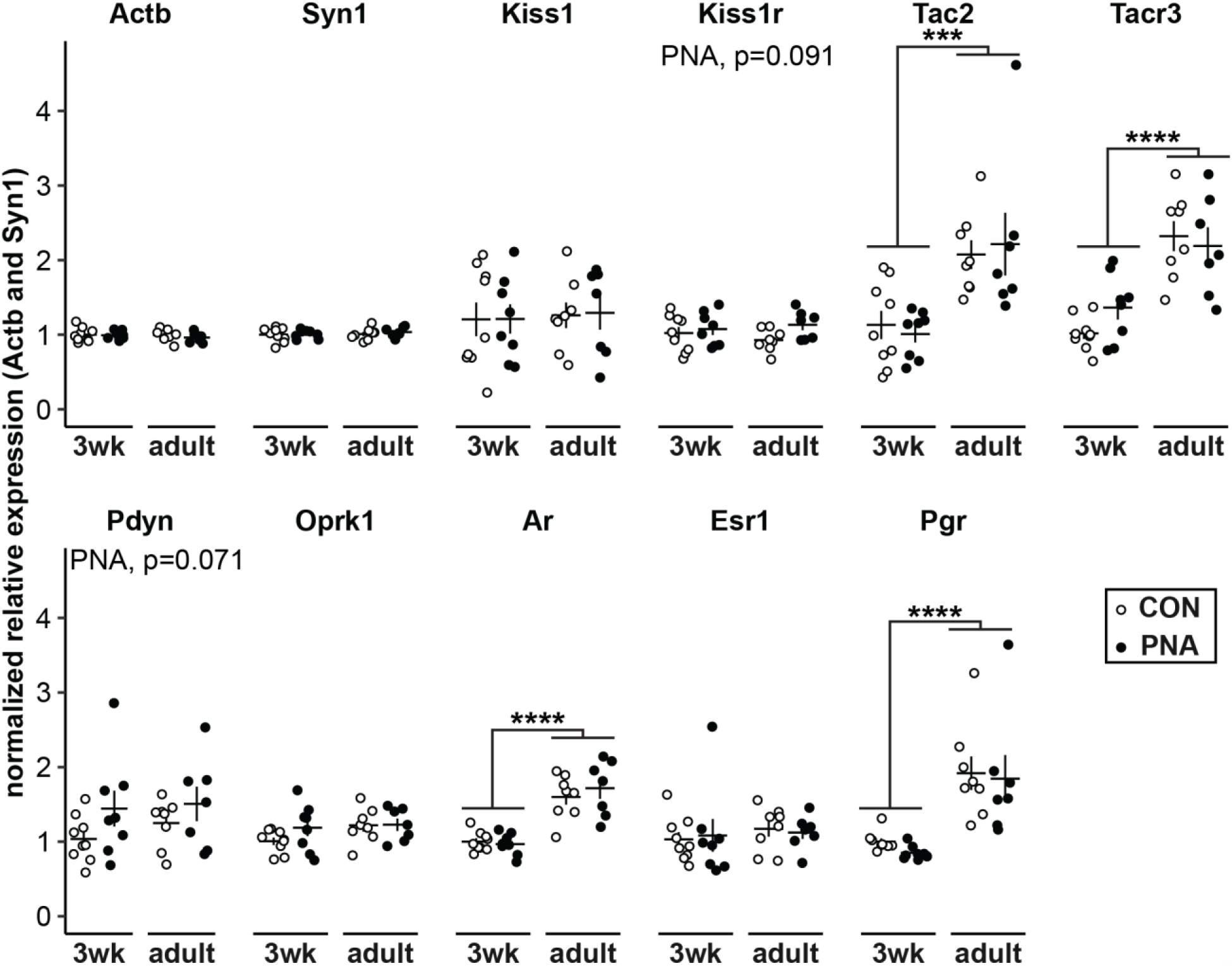
Effect of age and PNA treatment on arcuate nucleus mRNA transcripts. Individual values (CON open symbols, PNA black symbols) and mean ± SEM for *Actb, Syn1, Kiss1, Kiss1r, Tac2, Tacr3, Pdyn, Oprk1, Ar, Esr1*, and *Pgr* mRNA isolated from arcuate nucleus tissue punches. Statistical parameters are reported in Table 6. ***p≤0.001, ****p≤0.0001

## Discussion

Changes in the frequency of GnRH release throughout female reproductive cycles are important for fertility. Patients with PCOS often fail to exhibit these changes, instead displaying a persistently elevated LH, and presumably GnRH, pulse frequency. Here, we tested the hypothesis that prenatal exposure to elevated androgens, a model that recapitulates aspects of PCOS, disrupts the hypothalamo-pituitary-gonadal axis in part by changing the firing activity of KNDy neurons in the arcuate nucleus. Contrary to our hypothesis, neither overall spontaneous activity of KNDy neurons nor most burst characteristics were altered by PNA treatment either before puberty or in adulthood. Expression of *Tac2, Tacr3, Ar*, and *Pgr* mRNA was greater in the arcuate of adult mice as compared to 3wk mice, but this expression was not impacted by PNA treatment. These findings suggest that changes in KNDy neuron activity alone are not responsible for the altered LH pulse frequency observed with PNA treatment.

KNDy neurons have been postulated to be the pulse generator for GnRH release (Clarkson et al., 2017; McQuillan et al., 2019). Kisspeptin increases GnRH release (Glanowska et al., 2014; Messager et al., 2005) *in vivo* and in brain slices, and increases GnRH neuron activity (S.-K. Han et al., 2005; Pielecka-Fortuna et al., 2008) in brain slices. As the putative pulse generator, KNDy neuron activity would be expected to change in manners that reflect the output of GnRH neurons and LH release. GnRH neuron firing rate changes with development and PNA treatment alters the typical developmental trajectory. Specifically, in GnRH neurons from adults, firing rate is elevated in PNA mice (Roland and Moenter, 2011), whereas firing frequency in cells from PNA mice before puberty at 3wks of age is reduced, because PNA treatment blunts the typical peak in firing that occurs near this age in control mice (Dulka and Moenter, 2017).

We expected similar effects in KNDy neurons. Consistent with prior studies in adults, many of the KNDy cells that we recorded were quiescent (de Croft et al., 2012; Frazao et al., 2013; Ruka et al., 2013). Surprisingly, neither age nor PNA treatment altered the mean firing frequency of KNDy neurons. Similarly, effects on short-term burst firing were minimal. These observations suggest that both development and prenatal exposure to androgens alter GnRH neuron activity and release via mechanisms other than changing the activity of KNDy neurons.

The elevated LH pulse frequency in patients with PCOS is attributable at least in part to reduced negative feedback actions of progesterone (Pastor et al., 1998). When patients are treated with the anti-androgen flutamide, the suppressive effects of progesterone on LH release are partially restored (Eagleson et al., 2000), suggesting that hyperandrogenism plays a role in this impaired negative feedback. The opposing effects of androgens and progestins is supported by findings in murine brain slices that androgens interfere with progesterone negative feedback on GnRH neuron firing rate (Pielecka et al., 2006) and GABA transmission to these cells (Sullivan and Moenter, 2005). The elevated LH pulse frequency in PNA mice (Moore et al., 2015) may have a similar origin to that in patients with PCOS. Following ovariectomy, LH levels rise in control mice and to a lesser extent in PNA mice (Moore et al., 2015). Administration of progesterone reduces the LH levels in ovariectomized control but not PNA mice, indicative of impaired negative feedback in the latter (Moore et al., 2015). Progesterone may act in part through receptors in the arcuate nucleus to reduce LH pulse frequency, as administration of progesterone receptor antagonists in this brain region reduces the interval between LH pulses following intraperitoneal injection of progesterone (He et al., 2017). Progesterone also reduces the frequency of peaks in KNDy neuron activity measured by GCaMP fluorescence that are correlated with LH release (McQuillan et al., 2019), but whether or not this is a direct effect on KNDy neurons is not known.

A possible alternative mediator of central changes in PNA mice is GABAergic neurons. A subset of KNDy neurons may be GABAergic, though estimates vary on the percentage; up to 50% of KNDy neurons express GAD67 (Cravo et al., 2011), but only about 10-15% of KNDy neurons co-express the vesicular GABA transporter VGaT (Marshall et al., 2017). PNA treatment increases the frequency of GABAergic postsynaptic currents recorded in GnRH neurons from both adult and prepubertal 3wk-old mice (Berg et al., 2018; Sullivan and Moenter, 2004), which given the excitatory effects of GABA in GnRH neurons can contribute to increased activity. At least some of this increased transmission appears to arise from the arcuate nucleus as appositions between GABAergic neurons in the region and GnRH neurons increase in PNA mice (Moore et al., 2015). Consistent with an involvement in the steroid feedback effects discussed above, PNA treatment reduces the expression of progesterone receptors in GABAergic neurons in the arcuate nucleus (Moore et al., 2015). High-frequency optogenetic stimulation of GABAergic neurons in the arcuate can stimulate LH release in control mice (Silva et al., 2019). Longer-term chemogenetic activation of these GABAergic neurons also increases LH release, disrupts estrous cycles and decreases the number of corpora lutea (Silva et al., 2019). Together, these results point to GABAergic neurons in the arcuate as a potential steroid-sensitive mediator of PNA treatment on GnRH activity.

It is important to point out that our findings do not rule out a possible role of KNDy neurons in modulating the effects of PNA or age on GnRH and LH release. KNDy neurons are part of an intricate network in the arcuate nucleus, and they project to GnRH distal projections in the median eminence (Yip et al., 2015). In rats, PNA increases the number of arcuate cells immunopositive for kisspeptin and neurokinin B (Osuka et al., 2017) and the relative levels of kisspeptin and neurokinin B mRNA (Yan et al., 2014). In sheep, prenatal testosterone treatment reduced the number of putatively inhibitory dynorphin-immunopositive cells without changing the number of kisspeptin-immunopositive cells in the arcuate nucleus (Cheng et al., 2010). In contrast, following seven days of estradiol treatment, PNA mice did not differ in the relative mRNA expression of KNDy neuron peptides or receptors in the arcuate nucleus (Caldwell et al., 2015). Similarly, we only observed development changes in KNDy neuron peptide and receptor mRNA expression in the arcuate nucleus, but no changes due to PNA treatment. The variation between studies may be attributable to animal models or the examination of mRNA vs peptide. Prenatal testosterone exposure also alters synaptic connections of KNDy neurons with one another and projections to GnRH neurons (Cernea et al., 2015). It remains possible that even without a change in firing frequency, PNA may alter the amount of kisspeptin and/or other neuromodulators released at a given level of activity, potentially leading to increased GnRH release. The reciprocal connections of the network of KNDy neurons are also thought to be important for their involvement in GnRH pulse regulation, and PNA could alter these dynamics. Neurokinin B increases and dynorphin decreases the activity of KNDy neurons (de Croft et al., 2012, 2013; Ruka et al., 2013), whereas kisspeptin does not affect the firing frequency of other KNDy neurons (de Croft et al., 2013). The effects of neurokinin B and dynorphin signaling are modulated by steroidal milieu (Ruka et al., 2016). Intriguingly, in the present study, senktide, a neurokinin-3 receptor agonist, was less effective at eliciting an increase firing frequency of KNDy neurons from 3wk-old PNA mice. This suggests that PNA disrupts the development of this network.

A key feature of patients with PCOS is a persistently elevated LH pulse frequency that is most similar to the mid-to-late follicular phase (McCartney et al., 2002). Though we often recorded cells for at least 60min, these recordings were not of sufficient length to characterize rigorously the frequency and duration of peaks and nadirs in firing activity. It is plausible that the firing activity of KNDy neurons during a peak in activity does not differ with age or PNA treatment, but that the frequency of these peaks may be increased in adult PNA mice, leading to the elevated GnRH and LH pulse frequency. In this regard, the frequency of peaks, but not mean firing rate, was altered in KNDy neurons from adult males by orchidectomy and steroid replacement (Vanacker et al., 2017). Because PNA mice fail to exhibit typical estrous cycles and remain persistently in a diestrus-like state based on vaginal cytology, we specifically compared their firing frequency to that of cells from diestrous control mice. The frequency of peaks in calcium activity of KNDy neurons across the estrous cycle was not different from metestrus to diestrus to proestrus (McQuillan et al., 2019). In contrast, KNDy cells from mice in estrus exhibited a markedly decreased frequency of these peaks, postulated to be due in part to the negative feedback effects of progesterone (McQuillan et al., 2019). It is thus possible that a difference in firing rate of KNDy neurons would be observed on estrus that could be attributable to impaired progesterone negative feedback in adult PNA mice. Our choice to record on diestrus was based not only on the practical consideration that PNA mice are often in persistent diestrus, but also on the observed increase in LH pulse frequency in PNA mice during this stage (Moore et al., 2015). The lack of difference in KNDy neuron firing rate in the present study thus supports the postulate that this increased episodic LH release arises from other cells, or is disrupted by the brain slice preparation.

The work presented here indicates that the elevated GnRH firing frequency and LH pulse frequency associated with prenatal androgenization cannot be solely explained by changes in arcuate KNDy neuron firing frequency or bursting patterns. PNA may also alter the development of the stimulatory effects of neurokinin B receptor activation on KNDy neuron activity, disrupting network dynamics. Our work points to the importance of the broader network of neurons within the hypothalamus, including GABAergic cells, as mediators of the effects of hyperandrogenism on the output of the hypothalamic-pituitary-gonadal axis.

## Acknowledgements

Supported by National Institute of Health/Eunice Kennedy Shriver National Institute of Child Health and Human Development P50HD028934 and R01HD104345. AGG was supported by the National Defense Science and Engineering Graduate Fellowship Program. We thank Elizabeth Wagenmaker for expert technical assistance and Joshua Gibson for computational guidance

## Abbreviations

ACSF: artificial cerebral spinal fluid
ANOVA: analysis of variance
ARC: hypothalamic arcuate nucleus
DHT: dihydrotestosterone
FSH: follicle-stimulating hormone
GABA: gamma-aminobutyric acid
GFP: green fluorescent protein
GnRH: gonadotropin-releasing hormone
KNDy: neurons co-expressing kisspeptin, neurokinin B and dynorphin
LH: luteinizing hormone
PCOS: polycystic ovary syndrome
PNA: prenatally androgenized
PND: postnatal day
SK: senktide
Std resid: standardized residual

**Extended Data 1**. Code used to detect and analyze events in Igor Pro (coding-project) and for data analysis in R (PNA-KNDy).

## Notes

**Conflict of interest:** The authors declare no competing financial interests.

https://gitlab.com/um-mip/coding-project

https://github.com/gibson-amandag/PNA_KNDy

### Competing Interest Statement

The authors have declared no competing interest.

